# A microparticulate based formulation to protect therapeutic enzymes from proteolytic digestion: phenylalanine ammonia lyase as case study

**DOI:** 10.1101/638130

**Authors:** Irene Pereira de Sousa, Charlotte Gourmel, Olena Berkovska, Michael Burger, Jean-Christophe Leroux

## Abstract

Phenylketonuria is a genetic disorder affecting the metabolism of phenylalanine (phe) due to a deficiency in the enzyme phenylalanine hydroxylase. This disorder is characterized by an elevated phe blood level, which can lead to severe intellectual disabilities in newborns. The current strategy to prevent these devastating consequences is limited to a life-long phe-free diet, which implies major lifestyle changes and restrictions. Recently, an injectable enzyme replacement therapy, Pegvaliase, has been approved for treating phenylketonuria, but is associated with significant side-effects. In this study a phe-metabolizing system suitable for oral delivery is designed to overcome the need for daily injections. Active phenylalanine ammonia-lyase (PAL), an enzyme that catalyzes phe metabolism, is loaded into mesoporous silica microparticles (MSPs) with pore sizes ranging from 10 to 35 nm. The surface of the MSPs is lined with a semipermeable barrier to allow permeation of phe while blocking digestive enzymes that degrade PAL. The enzymatic activity can be partially preserved *in vitro* by coating the MSPs with poly(allylamine) and poly(acrylic acid)-bowman birk (protease inhibitor) conjugate. The carrier system presented herein may provide a general approach to overcome gastro-intestinal proteolytic digestion and to deliver active enzymes to the intestinal lumen for prolonged local action.

## 1. Introduction

Phenylketonuria (PKU) is a genetic disease that affects the metabolism of phenylalanine (phe). It is caused by a wide range of mutations in the gene encoding the enzyme phe hydroxylase, which in the presence of the cofactor tetrahydrobiopterin (BH4), catalyzes the conversion of phe into tyrosine. This enzyme deficiency leads to an accumulation of phe in the blood and the brain causing, if untreated, severe intellectual disabilities, developmental problems, psychiatric symptoms, microcephaly, motor deficits, autism and seizures.^[1]^ PKU is inherited in an autosomal recessive fashion with a prevalence of 1 in 10,000 in Europe, making it the most prevalent metabolic disorder caused by inborn error.^[2]^ The standard treatment for PKU is a phe-free diet to be initiated right after birth and maintained for life.^[2]^ This diet completely excludes protein-rich meals while allowing a restricted amount of low-protein food. Therefore, to reach the necessary daily intake of amino acids, medical food has to be provided, which has poor palatability. The ability of PKU patients to follow this demanding diet rigorously decreases with age, and becomes a significant burden during adolescence and adulthood.^[3, 4]^ To address the unmet medical needs of PKU patients, BioMarin Pharmaceutical has developed two therapeutic strategies, namely Kuvan and Pegvaliase. The first one, approved by FDA in 2007, is an oral formulation of sapropterin dihydrochloride, a synthetic derivative of BH4 that functions as a joint key in misfolded phe hydroxylase, restoring the activity of the enzyme. This drug is used for the treatment of BH4-responsive PKU, but phe-free diet is nevertheless necessary in the majority of cases.^[5-7]^ The second one, Pegvaliase (PALYNZIQ^™^), is an enzymatic replacement therapy that was approved by FDA in 2018.^[8]^ It is administered subcutaneously once-daily and consist of a conjugate of poly(ethylene glycol) (PEG) and a recombinant phenylalanine ammonia lyase (PAL), which catalyzes the conversion of phe to ammonia and trans-cinnamic acid.^[9, 10]^ In contrast to Kuvan, Pegvaliase is in principle effective in all PKU patients, but unfortunately, its administration is associated with several adverse effects such as injection site reactions, arthralgia, hypersensitivity reactions, dizziness, vomiting, anaphylactic shock, and the development of antibodies against PEG and PAL.^[8]^

In view of these limitations, developing a universal formulation capable of delivering an enzymatic replacement therapy to a larger patient population in a non-invasive fashion (e.g., oral administration) would greatly improve the disease management. Nevertheless, the oral delivery of enzymes implies one major challenge: the gastro-intestinal (GI) stability of the administered protein.^[11]^ Indeed, PAL is rapidly inactivated in the GI tract by acidic gastric pH and GI enzymes.^[12]^

To this end, Bourget et al. demonstrated that when PAL was encapsulated in cellulose semipermeable microcapsules it maintained 50% of its activity at pH 3.^[13]^ Even though no data were reported on the stability of this system in presence of intestinal proteases, the microcapsules (containing 5 IU PAL) induced an average phe blood level decrease of 35% in rats after 2 days of daily treatment.^[13, 14]^ PEGylated PAL was also tested in an oral formulation, given its higher *in vitro* stability, and it reduced phe plasma level by 40% in a PKU mouse model upon administration of 14 IU of enzyme.^[15]^ However, the formulation was only effective when repetitive doses were administered (every 2 h), indicating a low GI tract activity. Another study reported that the incorporation of PAL into a silica matrix protected the enzyme from intestinal proteases *in vitro*. Nevertheless, the administration of these particles in a PKU mouse model only marginally reduced phe plasma levels.^[15, 16]^

Designing a carrier that can protect PAL from the acidic gastric environment as well as from the proteases, predominately present in the intestinal environment, is essential for the formulation of an effective oral dosage form. Enteric capsules are a well-established approach that could be exploited to protect the formulation in the acidic environment of the stomach. Nonetheless, additional approaches should be sought to prevent the enzyme’s degradation in the intestine. Enzyme immobilization in an inorganic support, such as mesoporous silica particles (MSP), is a convenient strategy due to its low cost, high surface area, tunable particle and pore size.

The aim of this work was to take advantage of the above-mentioned strategies and design a phe-metabolizing system suitable for oral delivery to overcome the need for daily injections. To this end, PAL-loaded MSP were designed. The particles were coated with a protective semipermeable polymeric membrane shielding PAL from intestinal proteases, while permitting the diffusion of phe into the carrier for metabolism by the PAL cargo (**Figure 1**). As the particles could potentially be loaded in enteric capsules, the focus of this study was to prevent or slow down proteolytic degradation of PAL in the gastro-intestinal space.

**Figure 1.**
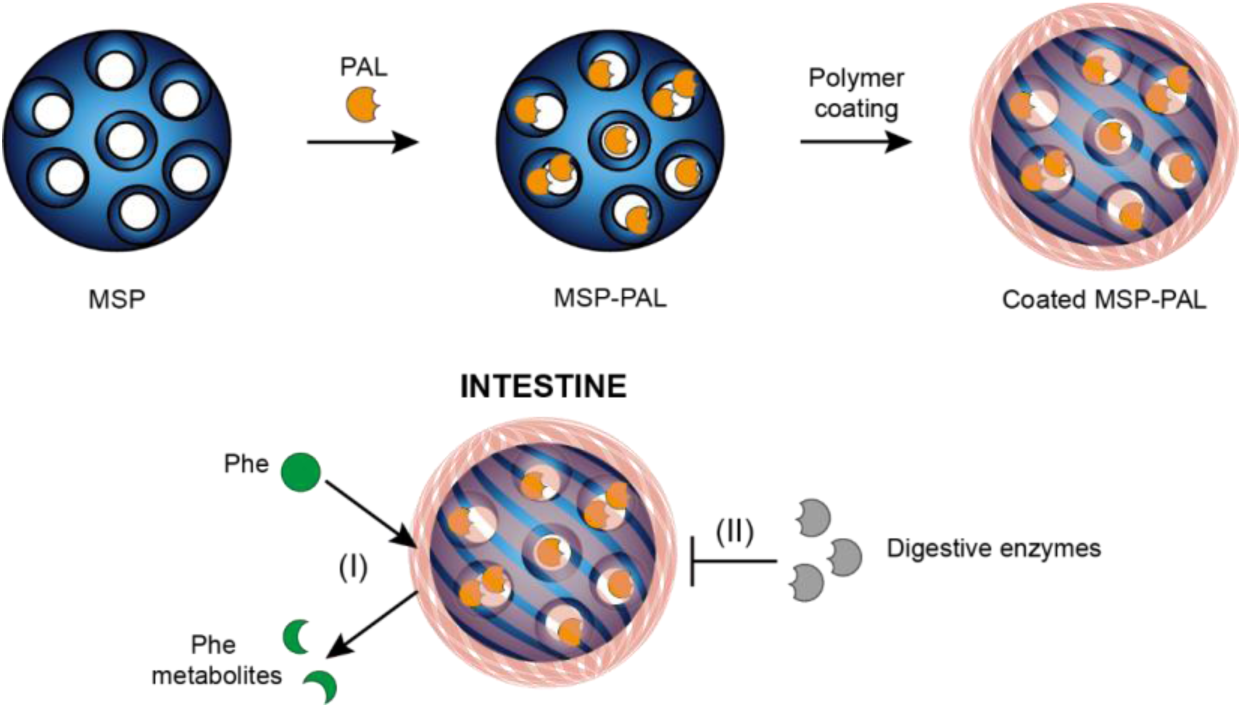
Schematic representation of the formulation, and of its proposed function in the intestinal tract: (I) diffusion of phe through the coating followed by its breakdown in the presence of loaded PAL and release of small-molecular phe metabolites from the formulation. (II) By blocking the diffusion of digestive enzymes, the polymer coating protects PAL against enzymatic degradation.

## 2. Results

### 2.1. Formulation and characterization of MSPs

MSPs with small pore sizes (MSP-s) were prepared by the template removal method using poloxamer P-123 (P-123) as the surfactant, tetraethylorthosilicate (TEOS) as the silica precursor, and mesytilene as the pore expander. Particles with larger pore sizes (MSP-l) were obtained following a hydrothermal treatment in a Teflon-lined autoclave.^[17]^ The template was efficiently removed, as confirmed by thermogravimetric analysis (TGA) (**Figure 2 a**). All formulations exhibited a mean particle size of approximately 12 – 20 μm. Pore diameter was on average 35 nm for MSP-l and 13 nm for MSP-s (**Figures 2 c-d, Table 1**). The expansion of the pore size in MSP-1 resulted in a slight decrease of the specific surface area in comparison to MSP-s (**Table 1**). In their dry state, the particles appeared as agglomerates of condensed beads with a porous surface, as shown by scanning electron microscopy (SEM) (**Figure 2 e**). To visualize the porous structure of MSPs, the particles were imaged by cryo-SEM after a freeze-drying step that enabled the removal of the first layers of water (**Figure 2 f**). Imaging of non-freeze-dried MSP-l by cryo-SEM, revealed the condensed-beads shape of the particles, but the surface topography could not be analyzed (**Figure S1**).

**Table 1.** Characterization of MSP-s and MSP-l in terms of specific surface area, pore volume, pore size, particle size, span and zeta potential. Data represent the mean ± SD (n = 3).

|  | MSP-s | MSP-l |
| --- | --- | --- |
| Specific surface area [m<sup>2</sup>/g] | 476.0 ± 56.5 | 338.5 ± 6.9 |
| Pore volume, adsorption [cm<sup>3</sup>/g] | 1.6 ± 0.6 | 2.9 ± 0.1 |
| Pore size, adsorption [nm] | 13.3 ± 3.7 | 34.6 ± 0.5 |
| Particle size [μm] | 13.0 ± 1.5 | 16.7 ± 2.6 |
| Span | 1.3 ± 0.4 | 1.0 ± 0.1 |
| Zeta potential [mV] | -32.5 ± 2.5 | -30.5 ± 2.2 |

**Figure 2.**
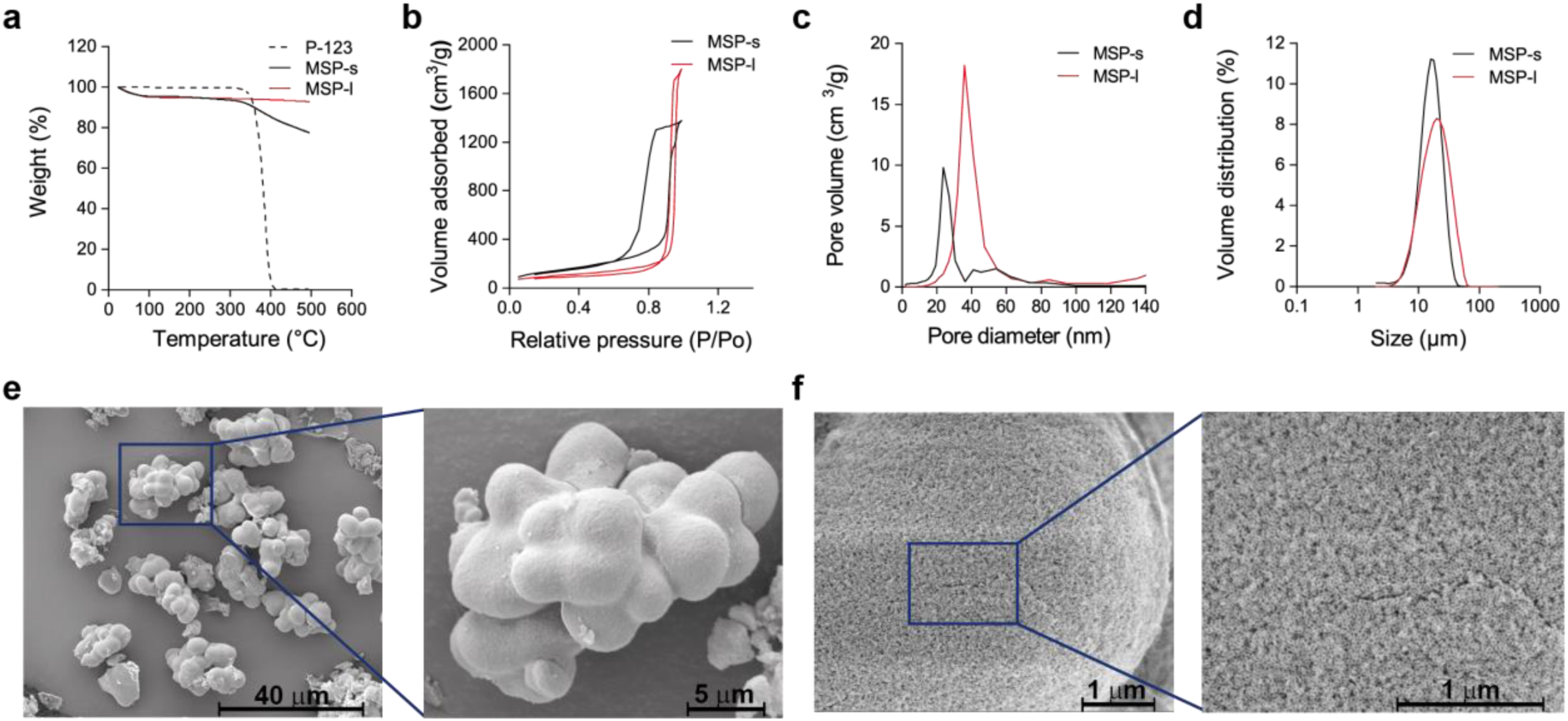
(a) TGA profiles of P-123, MSP-s and MSP-l. Nitrogen sorption isotherms (b), pore size distribution (c) and particle size distribution (d) of MSP-s and MSP-l. SEM (e, imaged with a voltage of 5 kV) and cryo-SEM (f, imaged with a voltage of 2 kV) images of MSP-l.

To introduce primary amino groups on the particles, MSP-l were functionalized with (3-aminopropyl)triethoxysilane (APTES) yielding a conjugation of 198 ± 13 µmol NH_2_/g particles (MSP-l-NH_2_). The functionalization did not impact the pore size (32 nm) nor the particle size (14 µm), which were comparable to non-functionalized MSP-l (**Figure S2 c - d, Table S1**). Furthermore, particle morphology was not affected by the functionalization. (**Figure S2 e - f**).

### 2.2. PAL loading into MSPs

The double mutant (C503S/C565S, Uniprot Q3M5Z3) PAL from *Anabaena variabilis* (AvPAL) in its PEGylated form is the active ingredient of Pegvaliase and was therefore selected for expression in this study.^[18]^ With our protocol an active protein (0.90 ± 0.20 IU/mg) was expressed, but with a low yield (0.65 ± 0.20 mg/L of culture). The expressed AvPAL was incorporated into MSP-s, MSP-l, and MSP-l-NH_2_ by adsorption.^[19]^ As reported in **Table S2**, the drug loading was higher for MSP presenting larger pores (8.5 ± 1.6% w/w) than for MSP with narrower pores (2.9 ± 1.8% w/w). Given its isoelectric point of 5.4, PAL is negatively charged at pH 7, the pH at which the loading step was performed.^[20]^ According to this, a greater entrapment efficiency would be expected for positively charged particles, compared to negatively charged ones.^[21]^ However, the incorporation of AvPAL in MSP-l-NH_2_ did not result in a significant increase of loading (+ 0.6% w/w), and therefore these particles were not further investigated.

As we could not produce AvPAL with sufficiently high yields to proceed with the evaluation of the formulations, the protein was substituted with a commercially available PAL (PAL from *Rhodotorula glutinis*, RgPAL). RgPAL is also a homotetrameric protein with a slightly higher subunit molecular mass (77 kDa) compared to AvPAL (64 kDa).^[20, 22]^ RgPAL was loaded into MSP-s to a similar extent as AvPAL (2.3 ± 0.3% w/w), while drug loading into MSP-1 was 30% lower with RgPAL in comparison to AvPAL (5.9 ± 0.5% w/w) (**Table S2**). These data confirmed the greater protein loading in the particles presenting larger pores, thus MSP-l were selected for further studies.

The characterization of the loaded and free RgPAL fractions by gel electrophoresis (**Figure S3 b**) revealed that the low-molecular weight contaminants of RgPAL were primarily incorporated into MSP, while a consistent fraction of the enzyme was found in the supernatant. Therefore, a separation method was set up yielding a purified PAL (pPAL) with similar secondary structure and kinetic parameters, but higher specific activity than RgPAL (**Figure S3 a - c**).

As reported in **Table 2**, the loading of pPAL into MSP-l was improved further from 5% to about 8% and finally 18% (w/w) by increasing the pPAL/MSP-l mass ratio from 1:10 to 2:10 and to 4:10, respectively. However, the loading resulted in a decrease in enzymatic activity, compared to free pPAL by ∼20%, 32%, and 56% for the 1:10, 2:10, and 4:10 ratios, respectively. A similar trend was observed when RgPAL was incorporated into MSP-l (**Table S3**).

**Table 2.** Encapsulation efficacy (ee), drug loading (dl), activity per mg pPAL, and maintained activity compared to free pPAL of pPAL in MSP-l at different PAL/MSP mass ratios. Data represent the mean ± SD (n = 3).

| PAL/MSP [w/w] | ee [%] | dl [%] | Activity [IU/mg<sub>pPAL</sub>] | Activity [%] |
| --- | --- | --- | --- | --- |
| 1:10 | 41.8 ± 8.9 | 4.8 ± 2.2 | 0.95 ± 0.33 | 79.9 ± 35.1 |
| 2:10 | 36.6 ± 9.0 | 7.9 ± 1.3 | 0.81 ± 0.10 | 67.8 ± 16.5 |
| 4:10 | 41.4 ± 2.5 | 18.5 ± 4.1 | 0.54 ± 0.16 | 43.8 ± 10.7 |

To confirm that pPAL was trapped inside MSPs and not only adsorbed to the surface, MSP-l were loaded with fluorescently labeled pPAL, and the particles were imaged by laser confocal scanning microscopy. The absence of free dye in the labeled pPAL mixture was confirmed by thin layer chromatography (TLC) (**Figure S4**). Z-stack images of MSP and fluorescently labeled MSP-PAL were collected, and the images of the central axial section of a particle both from the fluorescent channel, as well as from the bright field are shown in **Figure 3** (further representative images are reported in **Figure S5**). These images show that pPAL is present in the inner structure of MSP-l, as well as on the surface.

**Figure 3.**
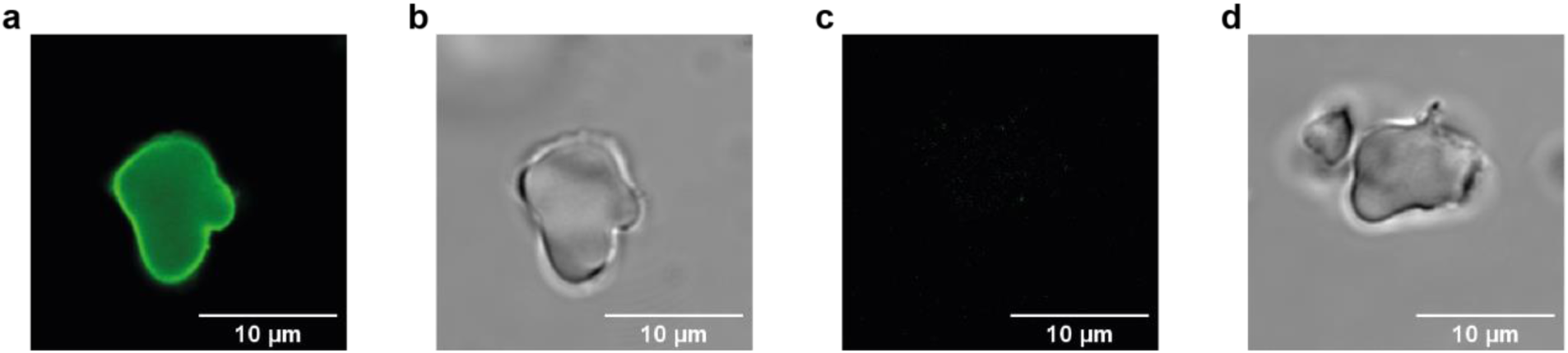
Representative confocal microscopy images of MSP-PAL loaded with fluorescent PAL in fluorescence channel (a) and in bright field (b), as well as of MSP in fluorescence channel (c) and bright field (d). Images have been collected by z-stacking and represent the central axial section of the particles.

In simulated intestinal fluids (SIF) (without proteases), the activity of pPAL was maintained for a longer period of time when encapsulated in MSP compared to the free enzyme, but no protection against proteases was provided by the particles resulting in a complete loss of PAL activity within 10 min (**Figure 4 c, Figure S6**).

**Figure 4.**
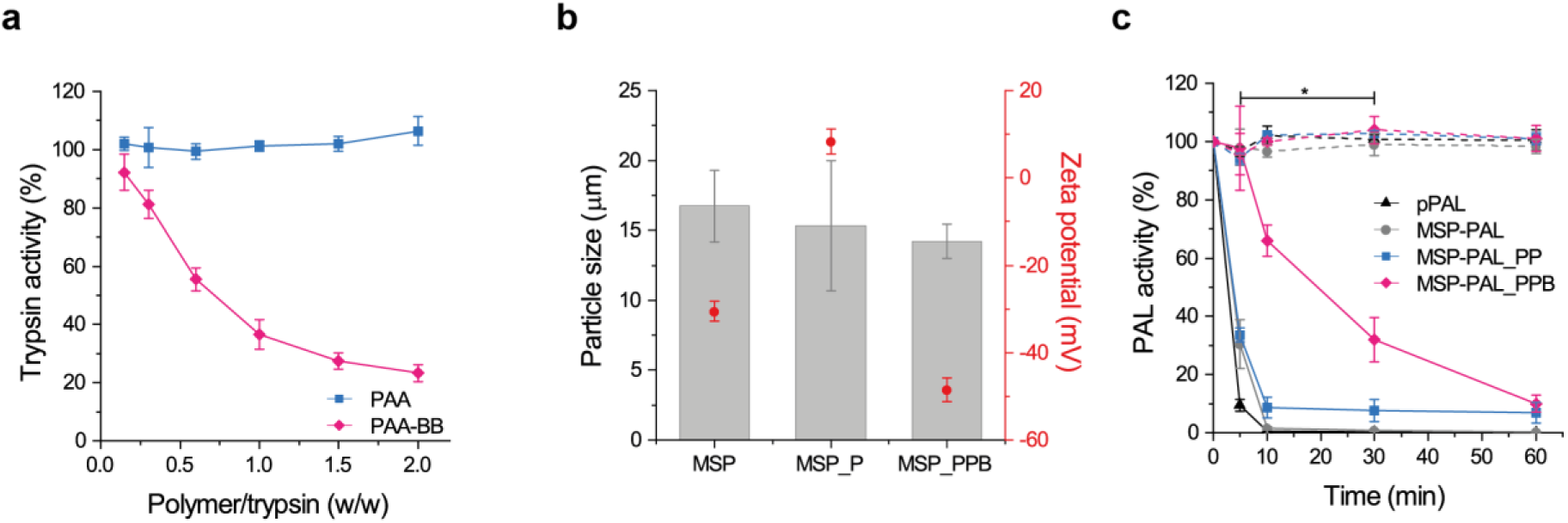
(a) Trypsin activity after 5 min incubation with increasing concentrations of PAA and PAA-BB. (b) Particle size and zeta potential of MSP, MSP coated with PAH (MSP_P) and MSP coated with PAH and PAA-BB (MSP_PPB). (c) Activity of pPAL, MSP-PAL, MSP-PAL coated with PAH and PAA (MSP-PAL_PP) and MSP-PAL_PPB in SIF (dotted lines) and in the presence of trypsin 0.4 mg/mL (solid lines) at 37 °C. Data represent the mean ± SD (n = 3), (c) two-ways ANOVA between trypsin treated MSP-PAL_PPB and trypsin treated MSP-PAL_PP (n = 5) (*p < 0.05).

### 2.3. Coating of MSP-PAL and protection against trypsin

To shield the encapsulated PAL from intestinal proteases, MSP-l were coated by the layer-by-layer technique starting with a first layer of a positively charged polymer, given the negative zeta potential of the particles (**Table 1**). Several polysaccharides and synthetic polymers were screened as coating materials, but they all showed poor protection against trypsin digestion (**Figure S7**). Therefore, to ameliorate the protection profile, a protease inhibitor was covalently linked to a polymeric backbone, and the conjugate was used as the coating material.^[23]^ It was previously reported that the protease inhibitor aprotinin was able to protect PAL from intestinal degradation *in vivo*.^[9]^ However, our screening showed that the inhibitor Bowman-Birk (BB) was more effective than aprotinin in protecting PAL from trypsin digestion, and thus was selected for further evaluation (**Figure S8** and **S9**). To have the protease inhibition in close proximity to the particles, BB was conjugated by carbodiimide mediated reaction to the negatively charged polymer poly(acrylic acid) (PAA), which was used as the second coating layer. Successful conjugation was confirmed by native gel electrophoresis performed at pH 4.1, which corresponds to the isoelectric point of BB (4.0 – 4.2)^[24]^. As illustrated in **Figure S10**, the BB conjugated to PAA migrated towards the positive pole, while free BB remained in proximity of the loading site. After conjugation, BB exhibited the same circular dichroism spectrum as the unconjugated protein (**Figure S11 a**), indicating that the conjugation did not result in an alteration of the protein structure. The polymer conjugate (PAA-BB) inhibited trypsin in a concentration-dependent fashion (**Figure 4 a**). By comparing the trypsin inhibition profile of PAA-BB and of free BB (**Figure S9 c**), it could be estimated that 0.15 mg of active BB per mg of PAA-BB was conjugated.

To follow the coating process, the particle size and zeta potential were determined after the deposition of the first coating layer (poly(allylamine) hydrochloride 15 kDa, positively charged) (PAH) and the second layer (PAA-BB, negatively charged). As represented in **Figure 4 b**, the particles maintained their original size during the process, while the zeta potential shifted from negative to positive, after the deposition of the first layer (MSP_P), and back to negative with the second layer (MSP_PPB). The ability of MSP_PPB (no PAL load) to inhibit trypsin was confirmed *in vitro* (**Figure S11 b**).

The double layer coating bearing BB protected the loaded pPAL from proteases, yielding a 35% PAL residual activity after 30 min incubation in the presence of trypsin. In the absence of the protease inhibitor (MSP-PAL_PP), only 8% activity remained at this time point. To confirm that the protective effect was provided by the conjugated BB and not by residual free BB possibly present in the PAA-BB mixture, the digestion assay was performed with MSP-PAL coated with PAH and then with a physical mixture of PAA and BB (PAA/BB) instead of the PAA-BB conjugate. These control particles did not produce a significant pPAL activity prolongation (**Figure S11 c**) confirming that the protective effect of MSP-PAL_PPB is due to the conjugated BB, and that traces of free BB, if any, are most likely eliminated to a large extent during particle purification.

## 3. Discussion

The management of the phe-free diet by PKU patient is extremely demanding especially during adolescence and adulthood. A lenient adherence to the treatment causes fluctuations in phe blood levels, which can result in severe consequences for the patients. Even though a new drug for PKU was recently approved, the treatment options are still scarce, leaving room for alternative therapies to address the unmet medical needs of this population.

The development of an oral enzyme replacement therapy would greatly improve the quality of life of PKU patients and facilitate the disease management for two reasons: (i) if the treatment is given in proximity of a meal it could degrade phe coming from the food, allowing the patient more flexibility with the diet; (ii) systemic phe is made accessible within the gut by its enterorecirculation, thus an oral enzymatic formulation could also degrade phe of systemic origin.^[25, 26]^ However, the design of efficient strategies to protect biologics from the intestinal environment still represents one of the biggest challenges in pharmaceutical sciences. Therefore, this work aimed at developing a carrier that could protect PAL from intestinal proteases while preserving its phe-metabolizing function.

A strategy to protect enzymes from GI proteolytic degradation is the conjugation to polymers that shield the enzyme from proteases.^[27]^ AvPAL conjugated with PEG from 5 to 20 kDa was previously tested for its stability *in vivo*, and only the enzyme conjugated with 5 kDa PEG showed a therapeutic relevant reduction of phe plasma levels, but repetitive doses (every 2 h) were required to maintain the effect.^[15]^ Even though other types of polymers/molecular architectures could be evaluated for conjugation (e.g., dendrimers ^[28]^), given the reported insufficient protection provided to PAL by polymer conjugation, we decided to pursue an alternative approach.

MSPs were selected as the enzyme carrier due to their tunable particle and pore size, and their ability to accommodate high-molecular weight proteins (such as PAL, ∼330 kDa ^[20]^) that maintain enzyme activity upon encapsulation to a large extent.^[29]^ MSP in the micrometer range (> 10 µm) were prepared to restrict particle intestinal uptake, for example by M cells, and thus guarantee a prolonged retention in the lumen while avoiding systemic exposure.^[30]^ Since the PAL diameter is 9.6 – 14.5 nm, a pore size greater than 15 nm would be desired to accommodate the enzyme.^[20]^ Previously, the encapsulation of PAL in a different type of MSP presenting a pore size ranging from 2 to 12 nm was reported.^[19]^ With this type of particles an enzyme loading of 5% was achieved, while with MSP-l, which have pore sizes of 35 nm, a PAL loading of up to 8% was obtained under comparable loading conditions (PAL/MSP mass ratio 2:10). Similarly to what was reported by Zhu et al.,^[19]^ while PAL encapsulation increased upon raising the PAL/MSP mass ratio, the activity decreased. This could be due to: (i) the inaccessibility of the substrate (phe) to the PAL located deeper in the pores; (ii) and/or macromolecular crowding, meaning that in pores presenting a high concentration of the enzyme, the mobility of the latter is reduced, which could impact its activity.^[29]^ However, the reduction of PAL activity was lower when encapsulated in MSP (−32%) compared to the encapsulation in silica particles (−69% ^[16]^) and in semipermeable microcapsules (−80% ^[13]^). To obtain a carrier with adequate loading of active PAL, MSP with the largest pore sizes (MSP-l, 35 nm) and a PAL/MSP mass ratio 2:10 were selected for this study.

Even though the MSP-l could accommodate PAL and prolong its stability in buffer compared to the free enzyme, these particles did not shield the enzyme from intestinal proteases (**Figure 4 c, Figure S6**). Therefore, a protective layer able to safeguard the loaded enzyme from intestinal proteases (e.g., trypsin 23.3 kDa), while allowing the permeation of the substrate (phe, 165 g/mol) through the coating was necessary to guarantee the function of the system. By exploiting the layer-by-layer technique, MSP-l could be coated with several combinations of positively and negatively charged polymers.^[31, 32]^ A negatively charged polymer was preferentially selected for the last layer (e.g., PAA) to provide a negative zeta potential, and thus reduce the interactions with the mucus layer, which is known to adhere and wrap around positively charged particles.^[33]^ The highest protective effect was achieved when a protease inhibitor-polymer conjugate was used for the coating. BB is an 8 kDa protein found in soybean and other mono- and dicotyledonous seeds that inhibits serine proteases.^[34]^ BB contains two independent binding sites for trypsin and for chymotrypsin, and is able to form binary (BB-trypsin or BB-chymotrypsin), as well as ternary (trypsin-BB-chymotrypsin) complexes with these proteases.^[24]^ The ability of BB to inhibit both proteases concomitantly makes it a very interesting agent to protect biomacromolecules from digestion in the intestinal tract. However, given the physiological function of proteases, their extensive inhibition in the lumen is undesirable as it could lead to undesired alteration of the digestive process.^[35]^ To prevent this, BB was covalently conjugated to the polymer used for the coating, allowing a localized inhibition of proteases that come in close proximity to the carrier. MSP-PAL particles coated with PAH and PAA-BB showed prolonged phe metabolization in the presence of trypsin *vs.* particles coated with PAH and PAA, indicating the fundamental contribution of BB in the inhibition of PAL digestion. The provided protection was lower than the one given by silica particles (50% maintained activity after 2 h in presence of trypsin ^[16]^). However, in this previous study^[16]^ the activity of the control silica particles (*i.e*., in absence of trypsin) was not stable during the experiment but, instead, increased as a function of time. This unusual aspect, which was not discussed in the manuscript, together with the fact that the activity is reported in absorbance, make the comparison with our data difficult. Even though our strategy provided a protective effect towards trypsin digestion, further optimizations would be possibly required to prolong PAL activity during the transit in the intestinal tract. To guarantee the gastric protection of the system the particle should be loaded into enteric capsules. However, testing these capsules *in vivo* would require a larger animal model than the gold standard Pah^enu2/enu2^ mouse model.

## 4. Conclusion

In this work, a phe-metabolizing system suitable for oral delivery was investigated. MSP with the large pore size (35 nm) proved to be an efficient carrier for high-molecular weight enzymes such as PAL, capable of loading high quantities of active enzyme. When encapsulated into MSP, PAL showed higher stability in buffer, however no protection against digestion by proteases was achieved. The key element to protect the loaded enzyme was the semipermeable coating composed of PAH and PAA-BB. Particularly, the conjugation of the protease inhibitor BB to PAA was necessary to provide protection, although further improvement of the system would be required before performing *in vivo* studies. It would be of interest to test the formulation loaded with PAL derivatives that are more resistant to protease digestion than the commercially available RgPAL. For instance, the company Codexis recently developed an AvPAL mutant more stable towards intestinal proteases (CDX-6114) and completed a Phase 1a clinical trial, with this enzyme, showing good tolerability at a dose up to 7.5 g (NCT03577886).^[36]^

Overall, given the tunability of the MSPs’ pore size and the possibility to coat the particles with a digestion-inhibiting polymeric layer, the proposed approach could provide a general mean to deliver orally various types of therapeutic enzymes and prolong their luminal activity.

## 5. Experimental Section

### Materials

Aprotinin, chitosan medium molecular weight, eppendorf LoBind tubes, kanamycin, Nα-benzoyl-L-arginine ethyl ester hydrochloride (BAEE), N-(3-dimethylaminopropyl)-N′-ethylcarbodiimide hydrochloride (EDAC), mesitylene, phe, RgPAL, PAH, P-123 (5.8 kDa), protease inhibitor cocktail, trifluoroacetic acid (TFA), silica gel 60 Å, TEOS, trypsin from porcine pancreas (13,671 IU/mg), trypsin-chymotrypsin inhibitor from glycine max soybean (bowman-birk, BB), and all salts and solvents were purchased from Sigma Aldrich (Buchs, Switzerland). The DNA fragment encoding for AvPAL was synthesized by Thermo Fischer GeneArt (Zug, Switzerland). The pET His6 TEV LIC cloning vector (1B) was a gift from Scott Gradia (Addgene plasmid # 29653; RRID:Addgene_29653). Dextran sulfate 40 kDa was purchased from Carl Roth (Karlsruhe, Germany), poly(acrylic acid) 50 kDa (PAA) was purchased from Polyscience Inc. (Warrington, PA, USA), and polystyrene sulfonate 70 kDa was purchased from ABCR GmbH (Karlsruhe, Germany). Pierce screw cap spin columns, micro BCA protein assay kit, BCA assay kit, T4 ligase (NEB, M0202), mouse HRP coupled anti-hexahistidine antibody and agarose UltraPure were purchased from ThermoScientific (Basel, Switzerland). 4,4-difluoro-5,7-dimethyl-4-bora-3a,4a-diaza-s-indacene-3-propionic acid succinimidyl ester (BDP FL NHS) fluorescent probe was obtained from Lumiprobe (Hannover, Germany) while APTES was bought from Acros Organics (Basel, Switzerland). Float-a-lyzer 50 kDa maximum sample volume 1 mL were purchased from Spectrum Labs (Breda, The Netherlands). PD MidiTrap G-25 column were obtained from GE Healthcare Life Sciences (Basel, Switzerland). Isopropyl β-D-thiogalactoside (IPTG) and lysozyme were purchased from Axon lab AG (Baden, Switzerland) while HindIII and Xhol restriction enzymes were purchased from BioConcept AG (Allschwil, Switzerland). Gel purification kit and Ni-NTA agarose was purchased from Qiagen (Hombrechtikon, Switzerland). DH5α competent *E.coli* and BL21(DE3)pLysS competent *E.coli* cells were purchased from Nimagen (Nijmegen, The Netherlands).

### Preparation of MSPs

MSPs were synthesized by adapting a previously described procedure.^[37]^ Briefly, 3 g of P-123 were dissolved in 90 mL of 2 M HCl at 40 °C. Then, 5.2 mL of mesitylene were added to the solution and stirred for 2 h (1000 rpm) at 40 °C. TEOS (6.82 mL) was added to the mixture and stirred (1000 rpm) for 14 h at 40 °C. To yield MSP-s, the suspension was heated at 120 °C for 10 h under reflux, without stirring. To yield MSP-l, the suspension was transferred to a Teflon vessel to be autoclaved at 170 °C for 5 h, and then allowed to cool down overnight. MSP-s and MSP-l were collected by centrifugation (4600 x *g*, 10 min). To remove the template, MSPs were dispersed in 250 mL of ethanol and stirred at room temperature for 2 h in a polypropylene vessel. The suspension was then sonicated for 15 min and filtered on paper. The solid was rinsed with ethanol and dried overnight at 80 °C.

### Preparation of MSP-l-NH_2_

The grafting of amine group to MSPs was achieved by adapting a previously described procedure.^[38]^ Briefly, MSP-l (0.4 g) were dried overnight at 80 °C under vacuum. The dried particles were dispersed in 65 mL of toluene, then, 0.4 mL of APTES were added. The mixture was refluxed at 80 °C for 6 h, under stirring. The functionalized particles (MSP-l-NH_2_) were collected by centrifugation (4600 x *g*, 15 min) and washed with 100 mL of ethanol. The particles were then dispersed in 30 mL of ethanol in a polypropylene vessel and sonicated for 15 min. Finally, the particles were filtered on paper and the solid was dried overnight at 80 °C.

To quantify the amount of amino groups present on MSPs a colorimetric assay was performed adapting a previously described protocol.^[39]^ Briefly, the primary amino groups present on MSP-l-NH_2_ (5 mg) were activated with 1 mL of 2-iminothiolane (20 × 10^-3^ M in bicarbonate buffer 0.1 M pH 8.5) for 1 h at room temperature. The sample was then centrifuged (18,000 x *g*, 5 min) and the pellet was washed twice with 1 mL water, twice with 1 mL ethanol, and twice with DTT 1 × 10^-3^ M to maintain the introduced sulfhydryl groups in the reduced state. The pellet was further washed twice with 1 mL ethanol and then twice with 1 mL of solution A (0.25 M bicarbonate buffer, pH 11.25). The washed pellet was dispersed in 1 mL of solution A and sonicated for 10 min in a water bath, 10 µL of this solution were withdrawn and added to 990 µL of BCA working solution (prepared following the provider’s instructions). The mixture was incubated for 1 h at 60 °C, then the absorbance was measured at 562 nm in a microplate reader (Infinite M200, Tecan, Männedorf Switzerland). The amount of primary amino groups was extrapolated from a calibration curve obtained by analyzing escalating concentrations of cysteine hydrochloride (0.05 – 0.8 × 10^-3^ M) under the same condition.

### Characterization of MSPs

To assess the complete extraction of the template, MSPs were analyzed by TGA (Q50, TA instruments, New Castle, DE, USA). MSPs were characterized for their size in ultrapure water (Mastersizer 2000, Malvern Panalytical, Malvern, UK) and for their zeta potential in phosphate buffer 4 × 10^-3^ M at a concentration of 0.4 mg/mL (ZetaView Z-NTA, Particle Metrix GmbH, Meerbusch, Germany). To assess the surface area and the pore diameter, 50 mg of dried MSPs were degassed for 1 h at 100 °C with nitrogen and then analyzed by nitrogen sorption at −196 °C (TriStar 3000, Micromeritics, Norcross, GA, USA). The morphology of MSPs was assessed by SEM (Quanta 200F, FEI, Eindhoven, Netherlands). For this analysis the dried particles were glued on a carbon support, the excess powder was eliminated with compressed air and the sample was sputter-coated with 5 nm Pt/Pd. The surface morphology of the samples was assessed by cryo-SEM (SEM LEO-1530, Zeiss, Oberkochen with cryo stage Leica, Vienna) at −120 °C after sample freeze fracture or after freeze fracture and freeze-drying performed as described in **Table S4**.

### Cloning, expression and purification of AvPAL

The synthesized DNA fragment encoding the full length AvPAL double mutant (C503S/C565S, Uniprot Q3M5Z3) was cloned into a modified version of the plasmid pET His6 TEV LIC (1B) at the restriction sites HindIII and XhoI (DNA sequence shown in **Table S5**). This resulted in the full length protein with a TEV protease cleavable hexahistidine tag at the N-terminus. Ligation was performed using T4 DNA ligase. The ligation product was transformed into DH5α cells and, subsequently, confirmed by sequencing (Microsynth AG, Balgach, Switzerland). The correct plasmid was then transformed into BL21(DE3)pLysS cells for protein expression. A small-scale expression test was performed by growing the transformed bacteria to an OD_600_ of 0.6 in LB medium supplemented with 50 µg/mL kanamycin and 30 µg/mL chloramphenicol. Protein expression was induced with IPTG 1 × 10^-3^ M and shaking at 250 rpm for 16 h at room temperature. The expression of AvPAL was assessed by western blot; the bacteria samples were lysed in laemmli buffer (2-mercaptoethanol 0.1% v/v, bromophenol blue 0.005% w/v, glycerol 10% v/v, sodium dodecyl sulphate 2% w/v, Tris-HCl 63 × 10^-3^ M pH 6.8), and separated by gel electrophoresis on a 12% SDS-PAGE. The proteins were then blotted on a PVDF membrane and immunostained with HRP coupled anti-hexahistidine antibody (**Figure S12 a**).

Preparative expression and purification of AvPAL was carried out as follows: One liter of supplemented LB medium (same as above) was inoculated 1:50 (v/v) from a saturated starter culture and grown at 37 °C until an OD_600_ of 0.6 was reached. Then, the expression was induced with IPTG 0.4 × 10^-3^ M at room temperature for 16 h. The cells were harvested by centrifugation (4600 x *g*, 20 min, 4 °C) and the pellet was resuspended in 20 mL of buffer 1 (Tris 25 × 10^-3^ M, NaCl 150 × 10^-3^ M, glycerol 10% v/v, pH 8) supplemented with 1 mg/mL lysozyme and 0.5% v/v protease inhibitor mix. The pellet was incubated on ice for 30 min and then sonicated (10 cycles of 15 s on, 15 s off, 60% amplitude on ice). The suspension was centrifuged (15,000 x *g*, 45 min, 4 °C), the supernatant was collected and supplemented with imidazole to reach a concentration of 10 × 10^-3^ M. The solution was loaded on a Ni-NTA column (200 µL resin volume) equilibrated with buffer 1 supplemented with imidazole 10 × 10^-3^ M. The column was washed with 20 mL of buffer 1 containing imidazole 10 × 10^-3^ M and, then, AvPAL was eluted with 1 mL of buffer 1 supplemented with imidazole 250 × 10^-3^ M. The collected protein (1 mL) was dialyzed overnight at 4 °C in a float-a-lyzer (50 kDa) against 100 mL of buffer 1 without imidazole. The presence of AvPAL in each expression step was assessed by denaturing SDS-PAGE (**Figure S12 b**).

The concentration of obtained AvPAL was quantified by microBCA following the provider’s instructions. The AvPAL activity was assessed by incubation with phe 15 × 10^-3^ M dissolved in Tris buffer 100 × 10^-3^ M (pH 8.5) at 37 °C. After 10 min, the reaction was stopped by addition of HCl 6 M and the absorbance of the produced trans-cinnamic acid (tCA, phe metabolite) was measured at 290 nm. The concentration of tCA was extrapolated from a calibration curve obtained by analyzing tCA solutions (6 - 200 × 10^-6^ M) with the same method.

### Purification of RgPAL

RgPAL was purified with a silica gel-loaded spin column (60 Å silica, 200 mg/column). The column was equilibrated with sodium phosphate buffer (PB) 50 × 10^-3^ M (pH 7), then 100 µL of RgPAL (4 mg/mL in PB) were loaded and incubated for 10 min. The purified protein (pPAL) was collected by centrifugation (10 s) into protein-low binding tubes. The column was used for a maximum of four times. The purity of the collected sample was assessed by denaturing SDS-PAGE, while the secondary structure was evaluated by circular dichroism (J-815, Jasco, Pfungstadt, Germany).

The concentration of pPAL was quantified by microBCA following the provider’s instructions. The specific activity of pPAL was assessed at 37 °C in Tris buffer 100 × 10^-3^ M (pH 8.5), containing phe 37.5 × 10^-3^ M (total reaction volume 200 µL). After 5 min, the reaction was stopped by adding 50 µL TFA 20% (v/v) in water and the amount of produced tCA was quantified by HPLC. The analysis was performed with a Hitachi HPLC system (VWR, Dietikon, Switzerland) equipped with a Chromaster 5160 pump, a 5260 autosampler and a UV/VIS detector. The samples were eluted with 50% v/v acetonitrile and 50% v/v water-TFA 0.1% v/v at 0.5 mL/min in a RP-C18 column (LiChroCART 250-4, 100, 5 µm) and detected at 280 nm. The concentration of produced tCA was extrapolated from a calibration curve obtained by analyzing tCA solutions (3 - 400 × 10^-6^ M) with the same method.

The activity of RgPAL and pPAL was evaluated in Tris buffer 100 × 10^-3^ M (pH 8.5) in the presence of escalating concentrations of phe (0.05 – 15 × 10^-3^ M) by monitoring the increment in absorbance at 290 nm with a microplate reader (Infinite M200). Kinetic parameters (K_m_ and V_max_) were extrapolated from the Eadie-Hofstee plot.^[40]^

### Encapsulation of PAL into MSPs

AvPAL, RgPAL and pPAL were encapsulated in MSP-s and MSP-l by adsorption. The enzyme was added to a MSPs suspension 3.3 mg/mL in PB 50 × 10^-3^ M (pH 7) at increasing PAL/MSP mass ratios (1:10, 2:10 and 4:10). The mixture was stirred for 2 h in protein-low binding tubes. The loaded particles were collected by centrifugation (10,000 x *g*, 3 min) and washed three times by centrifugation and re-suspension in PB. The amount of PAL present in the washing fractions was quantified by microBCA. These values were used to indirectly calculate drug loading and encapsulation efficiency. The activity of the encapsulated PAL (MSP-PAL) was assessed by HPLC as described in the section purification of RgPAL.

To visualize the loaded PAL by microscopy, the enzyme was labeled with a fluorescent probe as follows: pPAL (3.5 mg/mL) was dialyzed in a float-a-lyzer against 400 mL of sodium carbonate buffer (0.1 M, pH 8.5) overnight at 4 °C. To 1 mL of dialyzed pPAL, 5 µL of BDP FL NHS (10 mg/mL in anhydrous DMSO) were added and incubated for 12 h at 4 °C under shaking. The unconjugated dye was removed by size exclusion chromatography with a PD MidiTrap G-25 column. The absence of free dye was confirmed by TLC (water:methanol 1:9 v/v). The obtained fluorescent PAL was loaded into MSP-l following the procedure described above. After purification by centrifugation the particles were suspended in PB. An aliquot was loaded on a glass slide, covered with a coverslip and sealed with nail polish. The sample was imaged by confocal laser scanning microscopy (FluoView 300, Olympus, Center Valley, PA).

### Synthesis and characterization of PAA-BB

BB was covalently conjugated to PAA by EDAC mediated reaction. PAA (40 mg) was dissolved in 10 mL of water and the pH was adjusted to 5.5. EDAC (0.532 mg, 2.77 µmol) was added to the mixture and stirred for 30 min, keeping the pH at 5.5. Then, BB (22.2 mg, 2.77 µmol) was added dropwise and stirred for 3 h. The conjugated polymer (PAA-BB) was purified by dialysis (cut-off 25 kDa) against water for 3 days and then lyophilized.

The covalent conjugation of BB to PAA was confirmed by native gel electrophoresis, performed adapting a previously described procedure.^[41]^ Briefly, native agarose gels (0.5 % w/v) was prepared in hot acetate buffer (20 × 10^-3^ M, pH 4.1). Samples were loaded into the gel with loading buffer (glycerol 50% v/v, bromophenol blue 0.01% w/v) and the electrophoresis was carried out for 30 min at 50 mA in a horizontal gel electrophoresis apparatus (Sub-Cell GT Cell, BioRad, Cressier, Switzerland) using acetate buffer (as above) as the running buffer. The gel was stained for 30 min with an aqueous solution of acetic acid 10% (v/v), methanol 25% (v/v), Coomassie Blue 0.05% (w/v) and then destained with acetic acid 10% (v/v) and methanol 25% (v/v).

The ability of PAA-BB to inhibit trypsin was evaluated. Escalating doses of PAA-BB (7.5 - 100 µg/mL) were incubated with trypsin (2.1 × 10^-6^ M) in PB pH 7. After 5 min, a 10 µL aliquot was withdrawn and mixed with 150 µL BAEE (250 × 10^-6^ M). The BAEE absorbance was monitored at 253 nm in a microplate reader (Infinite M200) for 5 min. As control, the test was performed with trypsin and PAA (7.5 −100 µg/mL) or BB (1 to 20 µg/mL) instead of PAA-BB. The remaining trypsin activity was expressed as percentage compared to the activity of native trypsin assessed with the same method (100%).

### Coating of MSPs and characterization

After encapsulation of pPAL into MSP-l, the particles were washed twice with water and then dispersed in PB 4 × 10^-3^ M to reach a MSP concentration of 4 mg/mL. To 500 µL of this suspension, 350 µL of PAH (2 mg/mL in PB 1 × 10^-3^ M) were added dropwise under constant shaking. After 10 min the coated particles were collected by centrifugation (10,000 x *g*, 3 min), dispersed in 500 µL of PB 4 × 10^-3^ M and sonicated for 5 s in a bath sonicator at room temperature. The particles were coated with a second layer (PAA-BB or PAA 2 mg/mL in PB 1 × 10^-3^ M) following the same procedure. Finally, the particles were washed three times with water by subsequent centrifugation and resuspension.

The particles coated with PAH (MSP_P) or with PAH and PAA-BB (MSP_PPB) were characterized for their size and zeta potential as described in the section characterization of MSPs. The ability of MSP_PPB to inhibit trypsin was also evaluated. For this test 0.7 mg of particles were incubated with trypsin 17 × 10^-6^ M in SIF (potassium dihydrogen phosphate 50 × 10^-3^ M, pH 6.8, without pancreatin) for 5 min at 37 °C (total reaction volume 0.5 mL). A 100-µL aliquot was withdrawn and diluted with 700 µL PB. The remaining trypsin activity of the obtained solution was analyzed following the procedure described in the section synthesis and characterization of PAA-BB. As control the test was performed with MSP coated with PAH and PAA (MSP_PP).

PAL loaded MSP-l were also coated with one layer of chitosan, three layers of chitosan-dextran sulfate-chitosan, or with successive layers of PAH and polystyrene to form 4, 8, 10, and 15 layers following the same procedure described above but using water as solvent.

### Stability in simulated intestinal fluids (SIF) and in presence of trypsin

The stability of free and encapsulated pPAL was evaluated in SIF without pancreatin and in the presence of trypsin. Solutions containing 70 µg of pPAL, free or loaded, were prepared in SIF or in trypsin 17 × 10^-6^ M (total reaction volume 0.5 mL). The solutions were incubated at 37 °C under constant shaking. The pPAL stability in SIF or in the presence of trypsin was monitored for 4 and 1 h, respectively. At scheduled time points, 100 µL sample were withdrawn and the activity analyzed by HPLC following the procedure described in the section purification of RgPAL. The activity was expressed as percentage of the activity at time 0 h, which was set to 100%. To assess the ability of the protease inhibitors to protect pPAL from trypsin digestion, the same experiment was performed in the presence of escalating doses of aprotinin (30 – 310 × 10^-6^ M) and BB (12 - 250 × 10^-6^ M).

### Statistical analysis

Data are presented as mean±standard deviation (SD) of at least three independent experiments. GraphPad Prism software version 7 (GraphPad Softwares, La Jolla, CA, USA) was used to perform statistical analysis. Two-ways ANOVA followed by Bonferroni’s post hoc test was used for the analysis and p < 0.05 was considered statistically significant in all analyses.

## Acknowledgements

This work was supported by ETH Zurich Postdoctoral Fellowship and Marie Curie Actions for People COFUND program. The authors acknowledge support of the Scientific Center for Optical and Electron Microscopy ScopeM of the Swiss Federal Institute of Technology ETHZ, particularly Luiz Grafulha Morales (for SEM), Falk Lucas (for cryo-SEM) and Joachim Hehl (for fluorescence microscopy). The authors acknowledge Prof. Sotiris E. Pratsinis and Gian Nutal Schädli for their support with the nitrogen sorption measurements as well as Rout Saroj Kumar for his support with the circular dichroism measurements. CG acknowledges the Royal Society of Chemistry for a mobility grant. The authors declare no conflict of interest.

