## Supporting information for "A microparticulate based formulation to protect therapeutic enzymes from proteolytic digestion: phenylalanine ammonia lyase as case study"

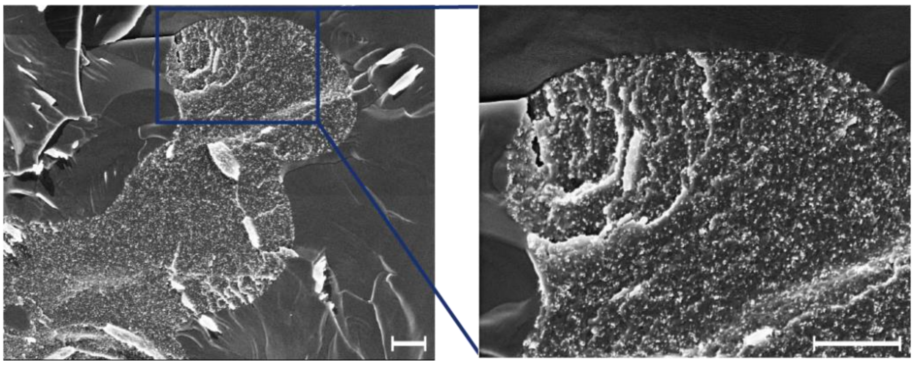

**Figure S1.** Cryo-SEM images of MSP-l without freeze-drying step. Scale bars represent 1 µm.

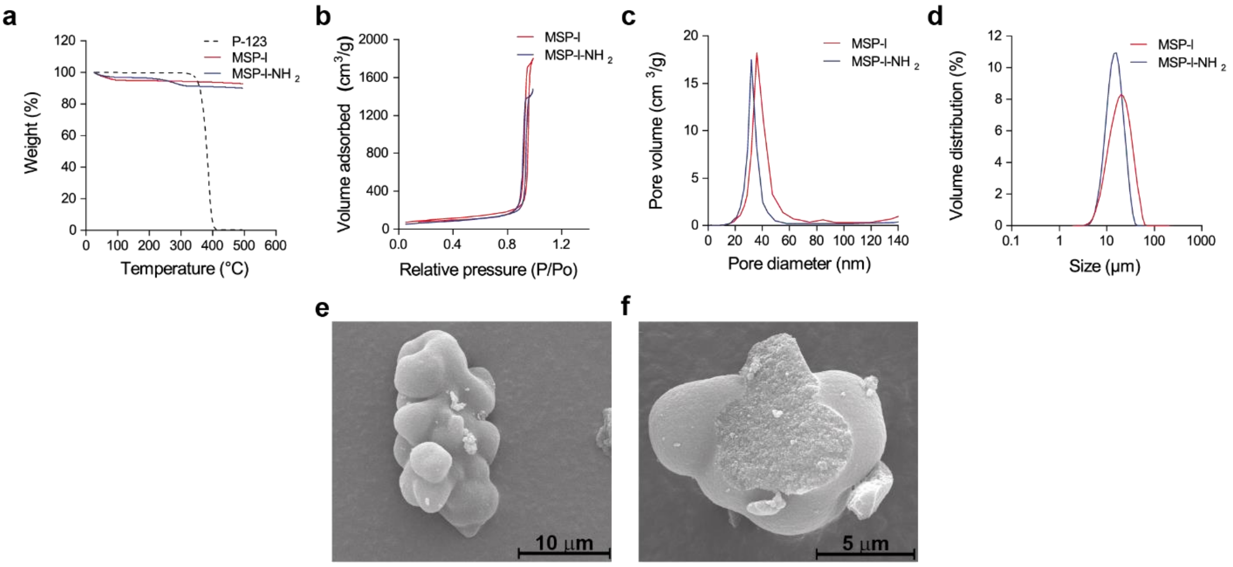


**Figure S2.** (a) TGA profiles of P-123, MSP-l and MSP-l-$NH_2$. Nitrogen sorption isotherms (b), pore size distribution (c), and particle size distribution (d) of MSP-l and MSP-l-$NH_2$. SEM images of an intact MSP-l-$NH_2$ (e) and a damaged particle exposing the porous inner structure (f).

a

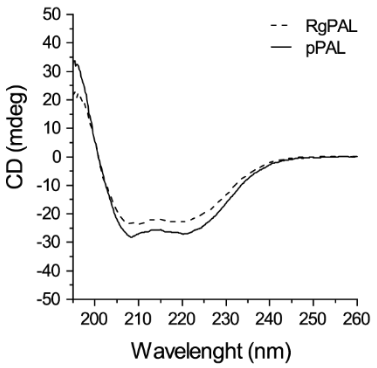


b

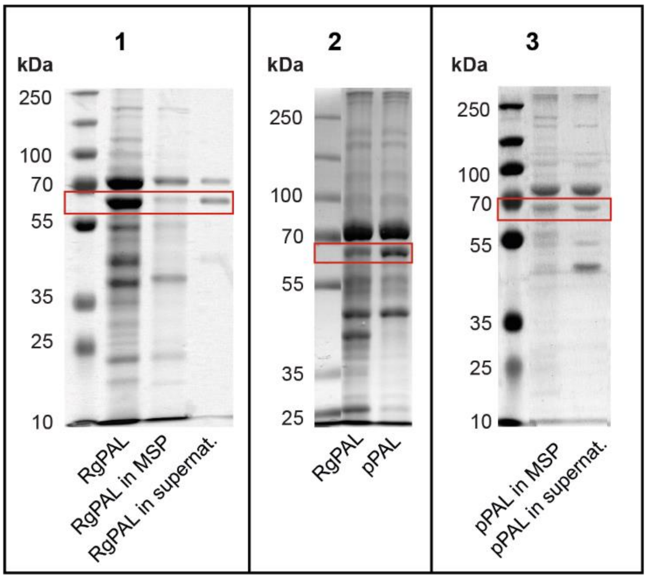


c

| | RgPAL | pPAL |
|---|---|---|
| $K_m$ (mM) | 0.61 ± 0.08 | 0.64 ± 0.11 |
| $V_{max}$ ( mM/min) | 0.019 ± 0.001 | 0.026 ± 0.01 |
| Specific activity (μmol/min/mg) | 0.74 ± 0.19 | 1.15 ± 0.25 |

**Figure S3.** (a) Circular dichroism spectra of PAL before (RgPAL) and after (pPAL) purification. (b1) SDS-PAGE of RgPAL, RgPAL loaded in MSP and of the free fraction. (b2) SDS-PAGE of RgPAL and pPAL. (b3) SDS-PAGE of pPAL loaded in MSP and of the free fraction. The area corresponding to the molecular weight of PAL's monomers is framed in red. (c) Comparison of activity and kinetic constants of RgPAL and pPAL.

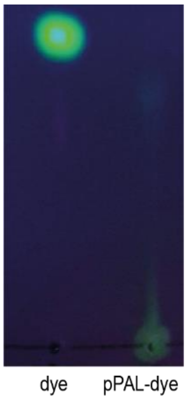


**Figure S4.** TLC of the dye (BDP FL NHS) and of the pPAL-dye conjugate.

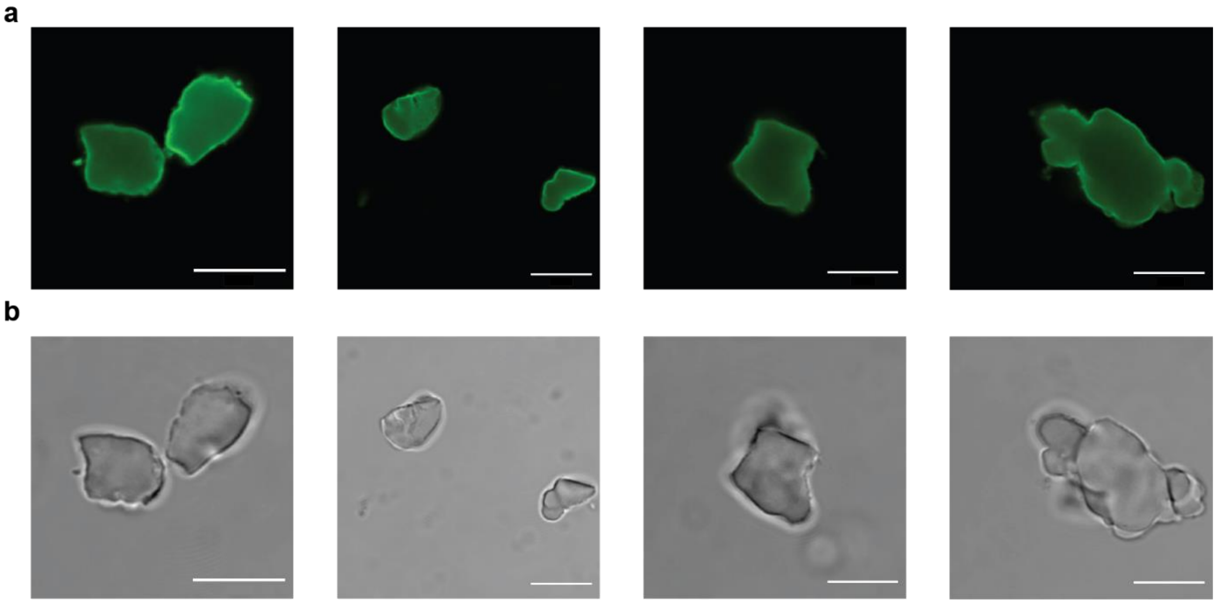


**Figure S5.** Additional representative confocal microscopy images of several MSP-PAL loaded with fluorescent PAL (fluorescence channel (a) and bright field (b)). Images have been collected by z-stacking and represent the central axial section of the particles. Scale bars represent 10 µm.

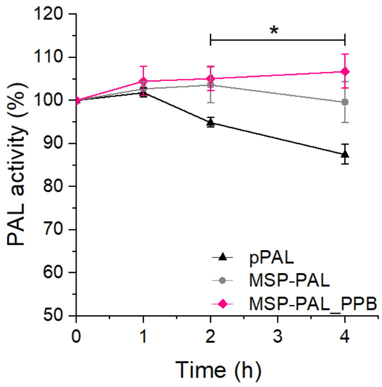


**Figure S6.** Activity of pPAL, MSP-PAL and MSP-PAL_PPB in SIF at 37 °C. Data represent the mean ± SD (n = 6), two-ways ANOVA between pPAL and MSP-PAL_PPB (*p < 0.05).

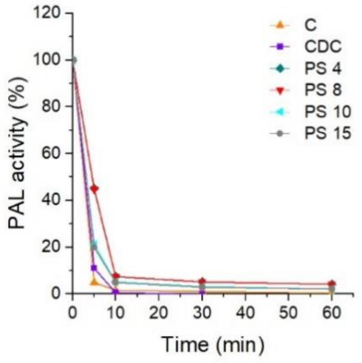


**Figure S7.** Activity of MSP-PAL coated with 1 layer of chitosan (C), chitosan-dextran sulfate-chitosan (CDC), PAH and polystyrene sulfonate in succession to form 4 layers (PS 4), 8 layers (PS 8), 10 layers (PS 10) and 15 layers (PS 15) in presence of trypsin 0.4 mg/mL.

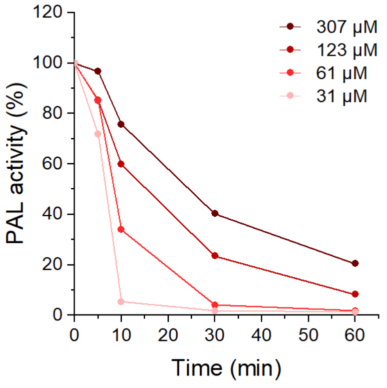


**Figure S8.** Activity of pPAL in presence of trypsin (0.4 mg/mL) and increasing concentrations of aprotinin (31, 61, 123, or 307 × $10^{-6}$ M).

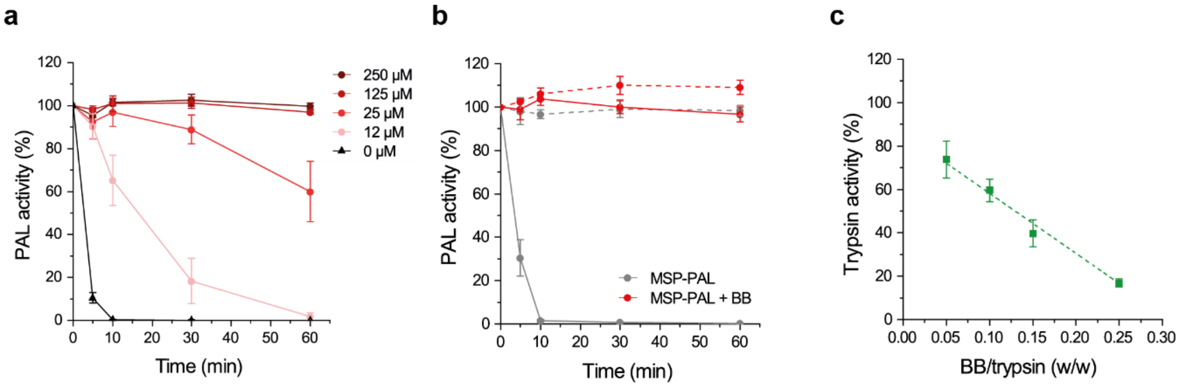


**Figure S9.** (a) Activity of pPAL in presence of trypsin (0.4 mg/mL) and increasing concentrations of BB (0, 12, 25, 125, 250 × $10^{-6}$ M). (b) Activity of MSP-PAL and MSP-PAL mixed with BB (125 × $10^{-6}$ M) in SIF (dotted lines) and in the presence of trypsin 0.4 mg/mL (solid lines). (c) Remaining trypsin activity after 5 min incubation with increasing concentrations of BB. Data represent the mean ± SD (n = 3).

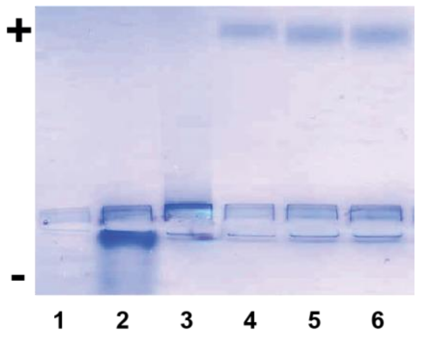


**Figure S10.** Agarose gel electrophoresis image of of PAA (1), BB (2), PAA/BB physical mixture (3), and PAA-BB from 3 independent syntheses (4, 5, 6) at pH 4.1 in acetate buffer.

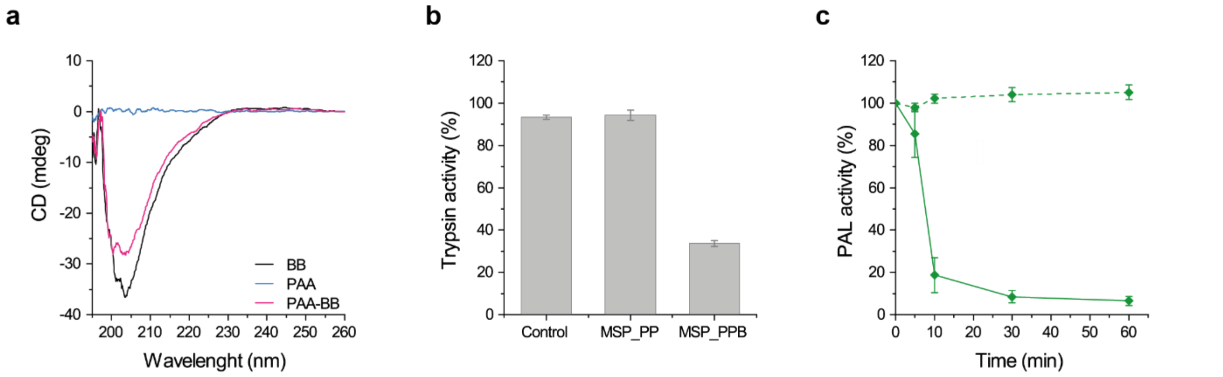


**Figure S11.** (a) Circular dichroism spectra of BB, PAA, and PAA-BB. (b) Trypsin activity after 5 min incubation with buffer (control), MSP coated with PAH and PAA (MSP_PP) and MSP coated with PAH and PAA-BB (MSP_PPB). (c) Activity of MSP-PAL coated with PAH and PAA/BB mixture in SIF (dotted lines) and in the presence of trypsin 0.4 mg/mL (solid lines). Data represent the mean ± SD (n = 3).

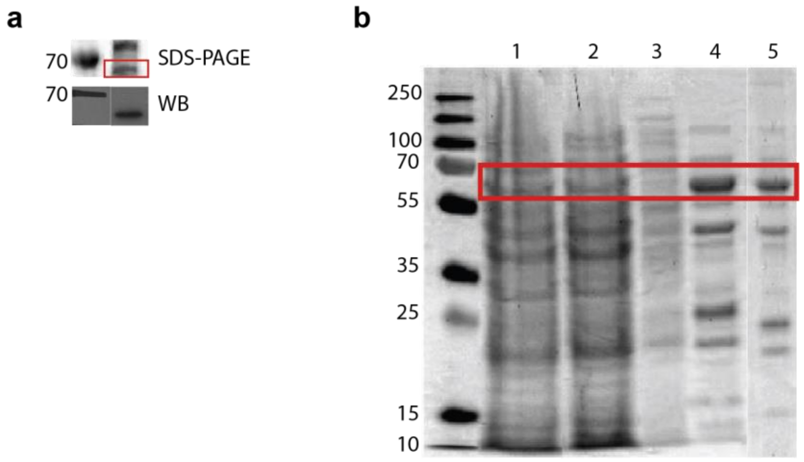


**Figure S12.** (a) SDS-PAGE and Western Blot (WB) stained for the hexahistidine tag of the pET His6 TEV LIC colony expressing AvPAL. (b) SDS-PAGE of the AvPAL expression steps: cell lysate supernatant (1), Ni-NTA flow-through (2), Ni-NTA wash with imidazole 10 × $10^{-3}$ M (3), elution with imidazole 250 × $10^{-3}$ M (4), AvPAL after dialysis (5). The area corresponding to the molecular weight of PAL's monomers is framed in red.

**Table S1.** Characterization of $MSP\text{-}l\text{-}NH_2$ in terms of specific surface area, pore volume, pore size, particle size, span and amount of conjugated primary amino groups.

| Characteristic | $MSP\text{-}l\text{-}NH_2$ |
|---|---|
| **Specific surface area [$m^2/g$]** | 284.8 |
| **Pore volume, adsorption [$cm^3/g$]** | 2.3 |
| **Pore size, adsorption [nm]** | 32.0 |
| **Particle size [µm]** | 13.8 |
| **Span** | 1.1 |
| **Amount of primary amino groups [µmol/g]** | 198 ± 13 |

**Table S2.** Encapsulation efficacy (ee) and drug loading (dl) of RgPAL or AvPAL in MSP-s, MSP-l, and $MSP\text{-}l\text{-}NH_2$ performed at a PAL/MSP mass ratio of 1:10. Data are expressed as mean ± SD (n = 3).

| MSP | ee [%] | dl [%] |
|---|---|---|
| **AvPAL/MSP-s** | 24.6 ± 12.6 | 2.9 ± 1.8 |
| **AvPAL/MSP-l** | 81.4 ± 3.5 | 8.5 ± 1.6 |
| **AvPAL/$MSP\text{-}l\text{-}NH_2$** | 86.9 ± 3.7 | 9.1 ± 1.8 |
| **RgPAL/MSP-s** | 18.6 ± 2.3 | 2.3 ± 0.3 |
| **RgPAL/MSP-l** | 54.7 ± 6.1 | 5.9 ± 0.5 |

**Table S3.** Encapsulation efficacy (ee), drug loading (dl), and activity per mg RgPAL of RgPAL in MSP-l at different PAL/MSP mass ratios. Data represent the mean ± SD (n = 3).

| PAL/MSP [w/w] | ee [%] | dl [%] | Activity [$IU/mg_{PAL}$] |
|---|---|---|---|
| **1:10** | 58.4 ± 3.4 | 5.9 ± 0.5 | 0.69 ± 0.11 |
| **2:10** | 63.4 ± 1.4 | 12.9 ± 1.5 | 0.44 ± 0.06 |
| **4:10** | 64.9 ± 5.1 | 26.3 ± 1.8 | 0.22 ± 0.04 |

**Table S4.** Freeze-drying steps for cryo-SEM sample preparation. The process was carried out in a BAF060 (Leica, Vienna) after freeze fracture. At the end of the process the samples were coated with tungsten *via* electron evaporation (5 nm, 45 ° angle with sample rotating at 40 rpm).

| Steps | $T_{start}$ [°C] | $T_{end}$ [°C] | Ramp [°C/h] | Hold at $T_{end}$ [h] |
|---|---|---|---|---|
| 1 | -120 | -115 | 30 | 1 |
| 2 | -115 | -110 | 30 | 1 |
| 3 | -110 | -100 | 30 | 1 |
| 4 | -100 | -90 | 30 | 1 |
| 5 | -90 | -80 | 30 | 1 |
| 6 | -80 | 20 | 30 | - |

**Table S5.** DNA sequence of the plasmid pET His6 TEV LIC (1B) containing the DNA fragment encoding for AvPAL. In the sequence are highlighted: T7 promoter (blue), starting methionine (red) hexahistidine tag (light blue) HindIII restriction site (green), TEV cleavage site (orange), AvPAL sequence (italic, double mutation in bold), stop codon (pink) and XhoI restriction site (violet).

| DNA sequence |
| --- |
| TGGCGAATGGGACGCGCCCTGTAGCGGCGCATTAAGCGCGGCGGGTGTGGTGGTTACGCGCAGCGTGA |
| CCGCTACACTTGCCAGCGCCCTAGCGCCCGCTCCTTTCGCTTTCTTCCCTTCCTTTCTCGCCACGTTCGC |
| CGGCTTTCCCCGTCAAGCTCTAAATCGGGGGCTCCCTTTAGGGTTCCGATTTAGTGCTTTACGGCACCTC |
| GACCCCAAAAAACTTGATTAGGGTGATGGTTCACGTAGTGGGCCATCGCCCTGATAGACGGTTTTTCGCC |
| CTTTGACGTTGGAGTCCACGTTCTTTAATAGTGGACTCTTGTTCCAAACTGGAACAACACTCAACCCTATCT |
| CGGTCTATTCTTTTGATTTATAAGGGATTTTGCCGATTTCGGCCTATTGGTTAAAAAATGAGCTGATTTAAC |
| AAAAATTTAACGCGAATTTTAACAAACTAGTAACGTTTACAATTTCAGGTGGCACTTTTCGGGGAAATGTGC |
| GCGGAACCCCTATTTGTTTATTTTTCTAAATACATTCAAATATGTATCCGCTCATGAATTAATTCTTAGAAAA |
| ACTCATCGAGCATCAAATGAAACTGCAATTTATTCATATCAGGATTATCAATACCATATTTTTGAAAAAGCC |
| GTTTCTGTAATGAAGGAGAAAACTCACCGAGGCAGTTCCATAGGATGGCAAGATCCTGGTATCGGTCTGC |
| GATTCCGACTCGTCCAACATCAATACAACCTATTAATTTCCCCTCGTCAAAAATAAGGTTATCAAGTGAGAA |
| ATCACCATGAGTGACGACTGAATCCGGTGAGAATGGCAAAAGTTTATGCATTTCTTTCCAGACTTGTTCAA |
| CAGGCCAGCCATTACGCTCGTCATCAAAATCACTCGCATCAACCAAACCGTTATTCATTCGTGATTGCGCC |
| TGAGCGAGACGAAATACGCGATCGCTGTTAAAAGGACAATTACAAACAGGAATCGAATGCAACCGGCGCA |
| GGAACACTGCCAGCGCATCAACAATGTTTTCACCTGAATCAGGATATTCTTCTAATACCTGGAATGCTGTT |
| TTCCCGGGGATCGCAGTGGTGAGTAACCATGCATCATCAGGAGTACGGATAAAATGCTTGATGGTCGGAA |
| GAGGCATAAATTCCGTCAGCCAGTTTAGTCTGACCATCTCATCTGTAACATCATTGGCAACGCTACCTTTG |
| CCATGTTTCAGAAACAACTCTGGCGCATCGGGCTTCCCATACAATCGATAGATTGTCGCACCTGATTGCCC |
| GACATTATCGCGAGCCCATTTATACCCATATAAATCAGCATCCATGTTGGAATTTAATCGCGGCCTAGAGC |
| AAGACGTTTCCCGTTGAATATGGCTCATAACACCCCTTGTATTACTGTTTATGTAAGCAGACAGTTTTATTG |
| TTCATGACCAAAATCCCTTAACGTGAGTTTTCGTTCCACTGAGCGTCAGACCCCGTAGAAAAGATCAAAGG |
| ATCTTCTTGAGATCCTTTTTTTCTGCGCGTAATCTGCTGCTTGCAAACAAAAAAACCACCGCTACCAGCGG |
| TGGTTTGTTTGCCGGATCAAGAGCTACCAACTCTTTTTCCGAAGGTAACTGGCTTCAGCAGAGCGCAGATA |
| CCAAATACTGTCCTTCTAGTGTAGCCGTAGTTAGGCCACCACTTCAAGAACTCTGTAGCACCGCCTACATA |
| CCTCGCTCTGCTAATCCTGTTACCAGTGGCTGCTGCCAGTGGCGATAAGTCGTGTCTTACCGGGTTGGAC |
| TCAAGACGATAGTTACCGGATAAGGCGCAGCGGTCGGGCTGAACGGGGGGTTCGTGCACACAGCCCAG |
| CTTGGAGCGAACGACCTACACCGAACTGAGATACCTACAGCGTGAGCTATGAGAAAGCGCCACGCTTCC |
| CGAAGGGAGAAAGGCGGACAGGTATCCGGTAAGCGGCAGGGTCGGAACAGGAGAGCGCACGAGGGAG |
| CTTCCAGGGGGAAACGCCTGGTATCTTTATAGTCCTGTCGGGTTTCGCCACCTCTGACTTGAGCGTCGAT |
| TTTTGTGATGCTCGTCAGGGGGGCGGAGCCTATGGAAAAACGCCAGCAACGCGGCCTTTTTACGGTTCCT |
| GGCCTTTTGCTGGCCTTTTGCTCACATGTTCTTTCCTGCGTTATCCCCTGATTCTGTGGATAACCGTATTAC |
| CGCCTTTGAGTGAGCTGATACCGCTCGCCGCAGCCGAACGACCGAGCGCAGCGAGTCAGTGAGCGAGG |
| AAGCGGAAGAGCGCCTGATGCGGTATTTTCTCCTTACGCATCTGTGCGGTATTTCACACCGCATATATGGT |
| GCACTCTCAGTACAATCTGCTCTGATGCCGCATAGTTAAGCCAGTATACACTCCGCTATCGCTACGTGACT |
| GGGTCATGGCTGCGCCCCGACACCCGCCAACACCCGCTGACGCGCCCTGACGGGCTTGTCTGCTCCCG |
| GCATCCGCTTACAGACAAGCTGTGACCGTCTCCGGGAGCTGCATGTGTCAGAGGTTTTCACCGTCATCAC |
| CGAAACGCGCGAGGCAGCTGCGGTAAAGCTCATCAGCGTGGTCGTGAAGCGATTCACAGATGTCTGCCT |
| GTTCATCCGCGTCCAGCTCGTTGAGTTTCTCCAGAAGCGTTAATGTCTGGCTTCTGATAAAGCGGGCCAT |
| GTTAAGGGCGGTTTTTTCCTGTTTGGTCACTGATGCCTCCGTGTAAGGGGGATTTCTGTTCATGGGGGTA |
| ATGATACCGATGAAACGAGAGAGGATGCTCACGATACGGGTTACTGATGATGAACATGCCCGGTTACTGG |
| AACGTTGTGAGGGTAAACAACTGGCGGTATGGATGCGGCGGGACCAGAGAAAAATCACTCAGGGTCAAT |
| GCCAGCGCTTCGTTAATACAGATGTAGGTGTTCCACAGGGTAGCCAGCAGCATCCTGCGATGCAGATCCG |
| GAACATAATGGTGCAGGGCGCTGACTTCCGCGTTTCCAGACTTTACGAAACACGGAAACCGAAGACCATT |
| CATGTTGTTGCTCAGGTCGCAGACGTTTTGCAGCAGCAGTCGCTTCACGTTCGCTCGCGTATCGGTGATT |
| CATTCTGCTAACCAGTAAGGCAACCCCGCCAGCCTAGCCGGGTCCTCAACGACAGGAGCACGATCATGC |
| GCACCCGTGGGGCCGCCATGCCGGCGATAATGGCCTGCTTCTCGCCGAAACGTTTGGTGGCGGGACCA |
| GTGACGAAGGCTTGAGCGAGGGCGTGCAAGATTCCGAATACCGCAAGCGACAGGCCGATCATCGTCGCG |
| CTCCAGCGAAAGCGGTCCTCGCCGAAAATGACCCAGAGCGCTGCCGGCACCTGTCCTACGAGTTGCATG |
| ATAAAGAAGACAGTCATAAGTGCGGCGACGATAGTCATGCCCCGCGCCCACCGGAAGGAGCTGACTGGG |
| TTGAAGGCTCTCAAGGGCATCGGTCGAGATCCCGGTGCCTAATGAGTGAGCTAACTTACATTAATTGCGT |
| TGCGCTCACTGCCCGCTTTCCAGTCGGGAAACCTGTCGTGCCAGCTGCATTAATGAATCGGCCAACGCG |
| CGGGGAGAGGCGGTTTGCGTATTGGGCGCCAGGGTGGTTTTTCTTTTCACCAGTGAGACGGGCAACAGC |
| TGATTGCCCTTCACCGCCTGGCCCTGAGAGAGTTGCAGCAAGCGGTCCACGCTGGTTTGCCCCAGCAGG |
| CGAAAATCCTGTTTGATGGTGGTTAACGGCGGGATATAACATGAGCTGTCTTCGGTATCGTCGTATCCCAC |
| TACCGAGATATCCGCACCAACGCGCAGCCCGGACTCGGTAATGGCGCGCATTGCGCCCAGCGCCATCTG |
| ATCGTTGGCAACCAGCATCGCAGTGGGAACGATGCCCTCATTCAGCATTTGCATGGTTTGTTGAAAACCG |
| GACATGGCACTCCAGTCGCCTTCCCGTTCCGCTATCGGCTGAATTTGATTGCGAGTGAGATATTTATGCCA |
| GCCAGCCAGACGCAGACGCGCCGAGACAGAACTTAATGGGCCCGCTAACAGCGCGATTTGCTGGTGACC |
| CAATGCGACCAGATGCTCCACGCCCAGTCGCGTACCGTCTTCATGGGAGAAAATAATACTGTTGATGGGT |

GTCTGGTCAGAGACATCAAGAAATAACGCCGGAACATTAGTGCAGGCAGCTTCCACAGCAATGGCATCCT
GGTCATCCAGCGGATAGTTAATGATCAGCCCACTGACGCGTTGCGCGAGAAGATTGTGCACCGCCGCTTT
ACAGGCTTCGACGCCGCTTCGTTCTACCATCGACACCACCACGCTGGCACCCAGTTGATCGGCGCGAGA
TTTAATCGCCGCGACAATTTGCGACGGCGCGTGCAGGGCCAGACTGGAGGTGGCAACGCCAATCAGCAA
CGACTGTTTGCCCGCCAGTTGTTGTGCCACGCGGTTGGGAATGTAATTCAGCTCCGCCATCGCCGCTTCC
ACTTTTTCCCGCGTTTTCGCAGAAACGTGGCTGGCCTGGTTCACCACGCGGGAAACGGTCTGATAAGAGA
CACCGGCATACTCTGCGACATCGTATAACGTTACTGGTTTCACATTCACCACCCTGAATTGACTCTCTTCC
GGGCGCTATCATGCCATACCGCGAAAGGTTTTGCGCCATTCGATGGTGTCCGGGATCTCGACGCTCTCC
CTTATGCGACTCCTGCATTAGGAAGCAGCCCAGTAGTAGGTTGAGGCCGTTGAGCACCGCCGCCGCAAG
GAATGGTGCATGCAAGGAGATGGCGCCCAACAGTCCCCCGGCCACGGGGCCTGCCACCATACCCACGC
CGAAACAAGCGCTCATGAGCCCGAAGTGGCGAGCCCGATCTTCCCCATCGGTGATGTCGGCGATATAGG
CGCCAGCAACCGCACCTGTGGCGCCGGTGATGCCGGCCACGATGCGTCCGGCGTAGAGGATCGAGATC
TCGATCCCGCGAAAT**TAATACGACTCACTATAGGG**GAATTGTGAGCGGATAACAATTCCCCTCTAGAAAT
AATTTTGTTTAACTTTAAGAAGGAGATATACC**ATG**GGTTCTTCT**CACCATCACCATCACCAT**ATTGATCCCT
TCACC**AAGCTT**GCGGATCCG**GAAAACCTGTATTTTCAG**_ATGAAAACCCTGAGCCAGGCACAGAGCAAAA_
_CCAGCAGCCAGCAGTTTAGCTTTACCGGCAATAGCAGCGCAAATGTGATTATTGGTAATCAGAAACTGAC_
_CATCAATGATGTTGCACGTGTTGCCCGTAATGGCACCCTGGTTAGCCTGACCAATAATACCGATATTCTGC_
_AGGGTATTCAGGCCAGCTGTGATTATATCAATAATGCAGTTGAAAGCGGTGAACCGATTTATGGTGTTACC_
_AGCGGTTTTGGTGGTATGGCAAATGTTGCAATTAGCCGTGAACAGGCAAGCGAACTGCAGACCAATCTGG_
_TTTGGTTTCTGAAAACCGGTGCAGGTAATAAACTGCCGCTGGCAGATGTTCGTGCAGCAATGCTGCTGCG_
_TGCAAATAGCCACATGCGTGGTGCAAGCGGTATTCGTCTGGAACTGATTAAACGCATGGAAATCTTTCTGA_
_ATGCCGGTGTTACCCCGTATGTTTATGAATTTGGTAGCATTGGTGCCAGCGGTGATCTGGTTCCGCTGAG_
_CTATATTACCGGTAGCCTGATTGGCCTGGACCCGAGCTTTAAAGTTGATTTTAATGGCAAAGAAATGGACG_
_CACCGACCGCACTGCGTCAGCTGAATCTGAGTCCGCTGACCCTGCTGCCGAAAGAAGGTCTGGCAATGA_
_TGAATGGCACCAGCGTTATGACCGGTATTGCAGCAAATTGTGTTTATGATACCCAGATTCTGACCGCAATT_
_GCAATGGGTGTTCATGCACTGGATATTCAGGCACTGAATGGTACAAATCAGAGCTTTCATCCGTTTATCCA_
_TAACAGCAAACCGCATCCGGGTCAGCTGTGGGCAGCAGATCAGATGATTAGCCTGCTGGCCAATAGCCA_
_GCTGGTTCGTGATGAACTGGATGGTAAACATGATTATCGTGATCATGAACTGATCCAGGATCGTTATAGCC_
_TGCGTTGTCTGCCGCAGTATCTGGGTCCGATTGTTGATGGTATTAGCCAGATTGCCAAACAAATCGAAATT_
_GAGATTAACAGCGTTACCGATAACCCGCTGATTGATGTTGATAATCAGGCAAGCTATCATGGTGGTAATTT_
_TCTGGGTCAGTATGTTGGTATGGGTATGGATCATCTGCGCTATTATATCGGTCTGCTGGCAAAACATCTGG_
_ATGTTCAGATTGCACTGCTGGCATCACCGGAATTTAGCAATGGTCTGCCTCCGAGTCTGCTGGGTAATCG_
_TGAACGTAAAGTTAATATGGGTCTGAAAGGTCTGCAGATTTGCGGTAATAGCATTATGCCGCTGCTGACCT_
_TTTATGGTAATAGTATTGCAGATCGTTTTCCGACCCATGCCGAACAGTTTAACCAGAATATTAACAGCCAG_
_GGTTATACCAGCGCAACCCTGGCACGTCGTAGCGTTGATATTTTTCAGAATTATGTTGCCATTGCCCTGAT_
_GTTTGGTGTTCAGGCAGTTGATCTGCGTACCTACAAAAAAACCGGTCATTATGATGCACGTGCC_***AGC***_CTG_
_TCACCGGCAACCGAACGTCTGTATAGCGCAGTTCGTCATGTTGTTGGTCAGAAACCGACCTCAGATCGTC_
_CGTATATTTGGAATGATAATGAACAGGGTCTGGATGAACATATTGCACGTATTAGTGCAGATATTGCAGCC_
_GGTGGTGTTATTGTTCAGGCCGTTCAGGACATTCTGCCG_***AGC***_CTGCAT_**TAATAA**AA**CTCGAG**CACCACCA
CCACCACCACTGAGATCCGGCTGCTAACAAAGCCCGAAAGGAAGCTGAGTTGGCTGCTGCCACCGCTGA
GCAATAACTAGCATAACCCCTTGGGGCCTCTAAACGGGTCTTGAGGGGTTTTTTGCTGAAAGGAGGAACT
ATATCCGGAT
